# Comparison of prokaryotic communities among fields exhibiting different disinfestation effects by anaerobic soil disinfestation

**DOI:** 10.1101/596825

**Authors:** Chol Gyu Lee, Toshiya Iida, Eriko Matsuda, Kayo Yoshida, Masato Kawabe, Masayuki Maeda, Yasunori Muramoto, Hideki Watanabe, Yoko Otani, Kazhiro Nakaho, Moriya Ohkuma

## Abstract

Anaerobic soil disinfestation (ASD) is a chemical-independent fumigation method used for reducing the abundance of pathogens at soil depths of <40 cm. However, its disinfestation efficiency is unstable under field conditions. The microbial community reflects the soil environment and is a good indicator of soil health. Therefore, soil with a good disinfestation efficiency may have a unique microbial community. The aim of the present study was to compare the prokaryotic communities among soils obtained from 17 geographically different greenhouses that experienced tomato bacterial wilt but exhibited different disinfestation efficiencies after ASD treatment with the same substrate. In the present study, soil prokaryotic communities in the field, which indicate difference in disinfestation effects after ASD treatment among several fields, were compared using next-generation sequencing. The prokaryotic communities in the fields showing different disinfestation effects were roughly separated into sampling fields. The relative abundances of *Betaproteobacteria* and *Clostridia* were significantly increased in well-disinfested fields. Overall, 25 operational taxonomic units (OTUs) were specifically increased in various well-disinfested soils and 18 OTUs belonged to phylogenetically diversified *Clostridia.* Other OTUs belonged to aerobic bacteria and were not previously detected in sample collected from ASD-treated fields. The results showed that the changes to the prokaryotic communities did not affect ASD efficiency, whereas changes in the abundance of specific microbes in the community were related to disinfestation.

## 1. Introduction

Soil-borne pathogens cause various plant diseases, including take-all, damping-off, crown rot, and wilting. Bacterial wilt caused by *Ralstonia solanacearum* has a host range exceeding 200 species from >50 families [1]. Soil disinfestation is challenging because this pathogen is distributed evenly at depths of >40 cm [2]. Several approaches have been attempted to control for bacterial wilt, including soil amendment, crop rotation, and field sanitation [3]. Although soil fumigation with chemical pesticides is an effective method for killing the pathogen causing bacterial wilt, the efficacy tends to be unstable in deep soil and the chemicals must escape to ensure food safety and prevent environmental pollution.

Anaerobic soil disinfestation (ASD) is an effective method to reduce the abundance of soil-borne pathogens [4]. This method comprises the incorporation of labile organic matter in the soil, irrigation, and covering the soil surface with polyethylene film. Organic matter increases microbial respiration, irrigation purges soil air, and polyethylene film prevents oxygen inflow from the atmosphere, which collectively induce reductive soil conditions [5,6]. Moreover, ASD using water-soluble organics, such as low-concentration ethanol or molasses as the carbon source, is effective for soil at depths of <40 cm [7]. Therefore, ASD using water-soluble carbon sources is suitable for the disinfection of *R. solanacearum* in deep-layer soils and is environmentally friendly. However, the disinfestation effects of ASD are unstable under field conditions [8]. A sufficient soil temperature, incubation period, and amount of carbon amendments are needed for the success of disinfestation.

Soil microbes reflect the soil environment and are considered an index of soil health [9,10]. ASD increases the abundance of several microbes that may be involved in the suppression of pathogens [11–16]. Therefore, fields with different disinfestation effects may have different soil microbial communities. Microbes that increase in abundance in well-disinfested soil that are commonly detected in several fields may be good candidate indicators for the efficiency of ASD treatment. The aim of the present study was to compare the prokaryotic communities among soils obtained from 17 geographically different greenhouses that had different disinfestation efficiencies after ASD treatment with the same substrate. The results of this multi-fields study showed that the soil microbial communities differed with the disinfestation efficiency of particular field and the well-disinfested fields had unique soil microbes, as compared with those not well-disinfested.

## 2. Materials and Methods

### 2.1. Sampling field and ASD treatment

Field experiments were established in 17 greenhouses situated on 8 fields in Japan that experienced bacterial wilt (Table 1).

**Table 1.** Characteristics of the sampled fields

| Field | Ishikawa | Hakusan | Niigata | Tsubame | Sanjo | Toyama | Gifu | Wakayama |
| --- | --- | --- | --- | --- | --- | --- | --- | --- |
| Code | Is | Ha | Ni | Ts | Sa | To | Gi | Wa |
| latitude and longitude | 36° 38' N, 136° 42' E | 36° 29' N, 136° 30' E | 37° 55' N, 139° 9' E | 37° 38' N, 138° 55' E | 37° 39' N, 139° 57' E | 36° 61' N, 136° 96' E | 35° 44' N, 136° 70' E | 33° 27' N, 135° 14' E |
| Soil types | Gleysol | Gleysol | Fluvisol | Fluvisol | Fluvisol | Gleysol | Gleysol | Cambisols |
| Replication | 1 | 3 | 3 | 3 | 2 | 3 | 1 | 1 |
| pH | 5.54 | 7.22 | 7.11 | 7.03 | 7.24 | 6.07 | 6.08 | 7.49 |
| EC (mS cm<sup>-1</sup>) | 0.03 | 0.31 | 0.32 | 0.24 | 0.20 | 0.29 | 0.12 | 0.88 |
| ASD conducted schedule | 7/27-8/18 | 7/11-7/28 | 8/18-9/9 | 7/31-8/21 | 9/1-9/22 | 8/2-8/26 | 8/11-9/1 | 7/29-8/19 |

Sugar-containing diatoms are discharged from food-processing facilities as by-products of the filtration of saccharified liquids. The main components of such by-products are sugars derived from the saccharified solution of tapioca starch and diatoms used as a filtering aid. These by-products, containing 40% by weight, were powdered and mixed into the soil with a rototiller at a ratio of 15 t ha^−1^ (approximately 6.0 g carbon kg soil^−1^) at a depth 30 cm. Thereafter, the field was covered with transparent polyethylene film (thickness, 0.1 mm) and flooded with more than approximately 150 L of water m^−2^. Each site was flooded at the time of disinfestation, and no irrigation was conducted afterward. Disinfestation was conducted for 21 days with the exception of field Ha (17 days) because the soil temperature of this field was >35°C during disinfestation. Each greenhouse (15 × 6 m) was subjected to ASD treatment. There were three replicates from fields Ha, Ni, Ts, and To; two from the field Sa; and none from fields Is, Gi, and Wa. Soil samples were collected from each greenhouse on the fields Is, Ha, To, Gi, and Wa before and after ASD treatment from two different depths—20–30 and 40–50 cm—using a core sampler (Gauge Auger DIK-106B; Daiki Rika Kogyo Co., Ltd, Saitama, Japan); i.e., 9 greenhouses × 2 depths × 2 sampling times = 36 soil samples in total. Only two soil samples (before and after ASD treatment in the upper layer soil) were collected from each greenhouse on fields Ni, Ts, and Sa (8 greenhouse × 2 sampling times = 16 soil samples). Overall, 52 soil samples were collected for analysis. Soil samples were collected from five randomly chosen points in each greenhouse and were mixed well.

### 2.2. Quantification of R. solanacearum in the field

The most probable number–polymerase chain reaction (MPN–PCR) method, which is a semi-quantitative *R. solanacearum* counting method [17], was conducted. Briefly, 10 g of soil was eluted into cultivation buffer and the soil extract was diluted with buffer to 10-, 100-, and 1000-fold. Each sample was incubated at 35°C for approximately 24 h. Thereafter, nested-PCR was performed using the samples as templates. The primer pair phcA2981f (5′-TGGATATCGGGCTGGCAA-3′) and phcA4741r (5′-CGCTTTTGCGCAAAGGGA-3′) was used in the first step of the PCR reaction and the primer pair phcA3538f (5′-GTGCCACAGCATGTTCAGG-3′) and phcA4209r (5′-CCTAAAGCGCTTGAGCTCG-3′) was used in the second step to target the *phcA*, which is key for the appearance of wilt disease. The number of samples that resulted in PCR products of ∼700-bp length was collectively checked against the MPN table. The detection limit of *R. solanacearum* using this method is 3–2400 colony forming units g^−1^ dry soil and a detection indicates the risk of outbreaks of bacterial wilt in the field.

DNA was extracted from 0.5 g of soil using the ISOIL for bead beating kit (Nippon Gene Co., Ltd., Tokyo, Japan), according to the manufacturer’s instructions. DNA quantification and integrity were measured using a NanoDrop spectrophotometer (Thermo Fisher Scientific, Waltham, MA, USA) and gel visualization (0.8% agarose in Tris/acetic acid/ethylenediaminetetraacetic acid buffer), respectively. The V4 region of the 16S rRNA gene of each sample was amplified by PCR using the bacterial and archaeal universal primers 515F (5′-GTGCCAGCMGCCGCGGTAA-3′) and 806R (5′-GGAC-TACVSGGGTATCTAA-3′) [18]. A library was prepared by adaptor ligation with the PCR primer pairs using the TruSeq Nano DNA Library Prep Kit (Illumina, Inc., San Diego, CA, USA). When two or more bands were detected using 1.5%-agarose gel electrophoresis, PCR products of approximately 300 bp in length were excised from the gel, non-specific amplicons were removed, and the products were purified using a MonoFas DNA purification kit for prokaryotes (GL Sciences, Inc., Tokyo, Japan). One soil sample (the upper layer of soil of To3 after ASD treatment) was not amplified after PCR amplification and was excluded from analysis. Each PCR amplicon was cleaned twice to remove the primers and short DNA fragments using the Agencourt AMPure XP system (Beckman Coulter, Inc., Brea, CA, USA) and quantified using a Qubit Fluorometer (Invitrogen Corporation, Carlsbad, CA, USA). Following successful amplification, the PCR products were adjusted to equimolar concentrations and subjected to unidirectional pyrosequencing, which was performed by Bioengineering Lab. Co., Ltd. (Kanagawa, Japan) using a MiSeq instrument (Illumina, Inc.). Overall, 3,114,333 sequences were obtained from the 51 samples (Supplemental Table 1). Sequencing data were deposited in the DNA Database of Japan Sequence Read Archive under the accession number DRA006673.

### 2.4. Data analyses

The sequencing data were analyzed as previously described [19]. Raw FASTA files were pre-processed with the Quantitative Insights into Microbial Ecology (QIIME) bioinformatics pipeline [20]. Data from the read sequences, quality, flows, and ancillary metadata were analyzed using the QIIME pipeline. Quality filtering consisted of discarding reads of <200 or >1000 bp in length, excluding homopolymer runs of >6 bp and continuous ambiguous bases of >6 bp, and accepting one barcode correction and two primer mismatches. Moreover, reads with a mean quality score of <25 were removed. Finally, singleton operational taxonomic units (OTUs) and chimeric sequences were removed for statistical analyses. Denoising was performed using the built-in denoiser algorithm, and chimera removal and OTU picking were accomplished using the USEARCH 61 sequence analysis tool (http://www.drive5.com/usearch/download.html) and a pairwise identity percentage of 0.97. Taxonomy assignment was performed using the Ribosomal Database Project naïve Bayesian classifier with a minimum confidence of 0.8 against the Greengenes database (October 2012 release; http://greengenes.secondgenome.com/) and the Basic Local Alignment Search Tool (https://blast.ncbi.nlm.nih.gov/Blast.cgi). The richness and diversity of pyrotag-based datasets were determined by OTU-based analysis using the phyloseq R package, version 1.7.24 [21]. The alpha diversity within each individual sample was estimated using the non-parametric Shannon diversity index. The Chao1 estimator was used to estimate the richness of each sample. A multivariate analysis of the community structure and diversity of the pyrotag-based datasets was performed using a weighted UniFrac dissimilarity matrix calculated in the QIIME, jackknifing (1000 reiterations) read abundance data at the deepest possible level (41,012 reads), and using unconstrained ordination by a principal coordinate analysis (PCoA) for prokaryotes in each soil layer. *K*-means clustering was used to verify the effects of the sampling fields and ASD treatment on prokaryotic communities. Finally, an indicator value analysis was conducted using the indicspecies R package [22] to identify OTUs associated with suppressive soil rather than conducive soil or vice versa (*p* < 0.05). Two soil group patterns were created. One group was divided into samples from the soil before and after ASD treatment. The other group was divided based on disinfestation efficiency. The datasets were merged and the abundant OTUs that were associated with the pathogen-free soil after ASD treatment were evaluated in each layer (*p* < 0.001). The number of random permutation tests for the calculation of the indicator values was 999.

### 2.5. Construction of phylogenetic tree

A phylogenetic tree of clostridial members was constructed by a 1000-fold bootstrap analysis using the neighbor-joining method with the Clustal W multiple sequence alignment algorithm (version 2.0) [23] and NJ plot software [24]. The phylogenetic tree contained the 16S rRNA sequences of several clusters of clostridial strains.

## 3. Results

### 3.1. Effectiveness of ASD treatment with sugar-containing diatoms

*R. solanacearum* in soil was not detected after ASD treatment in in the upper layers of 14 (82.4%) of 17 greenhouses and lower layers of 7 (77.8%) of 9 greenhouses (Table 2). Therefore, ASD treatment was appropriate in each field. These soils were defined as well-disinfested (disinfestation success soil: S-soil). However, some fields (Is, Ha1, and Ha2) were not well disinfested (disinfestation failure soil: F-soil).

**Table 2.**
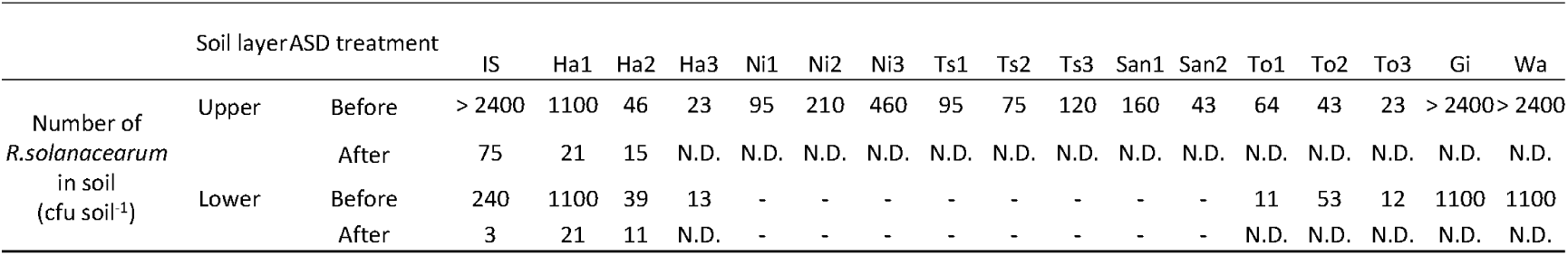
The number of *R. solanacearum* in each field before and after ASD treatment

### 3.2. Changes of prokaryotic communities after ASD treatment

The sequences were clustered into 74,539 OTUs at 97% similarity. Among the S- and F-soils, the DNA concentrations were significantly decreased in the upper layer of the S-soils and the lower layer of the F-soils (Figure 1, Supplemental Table 2).

**Figure 1.**
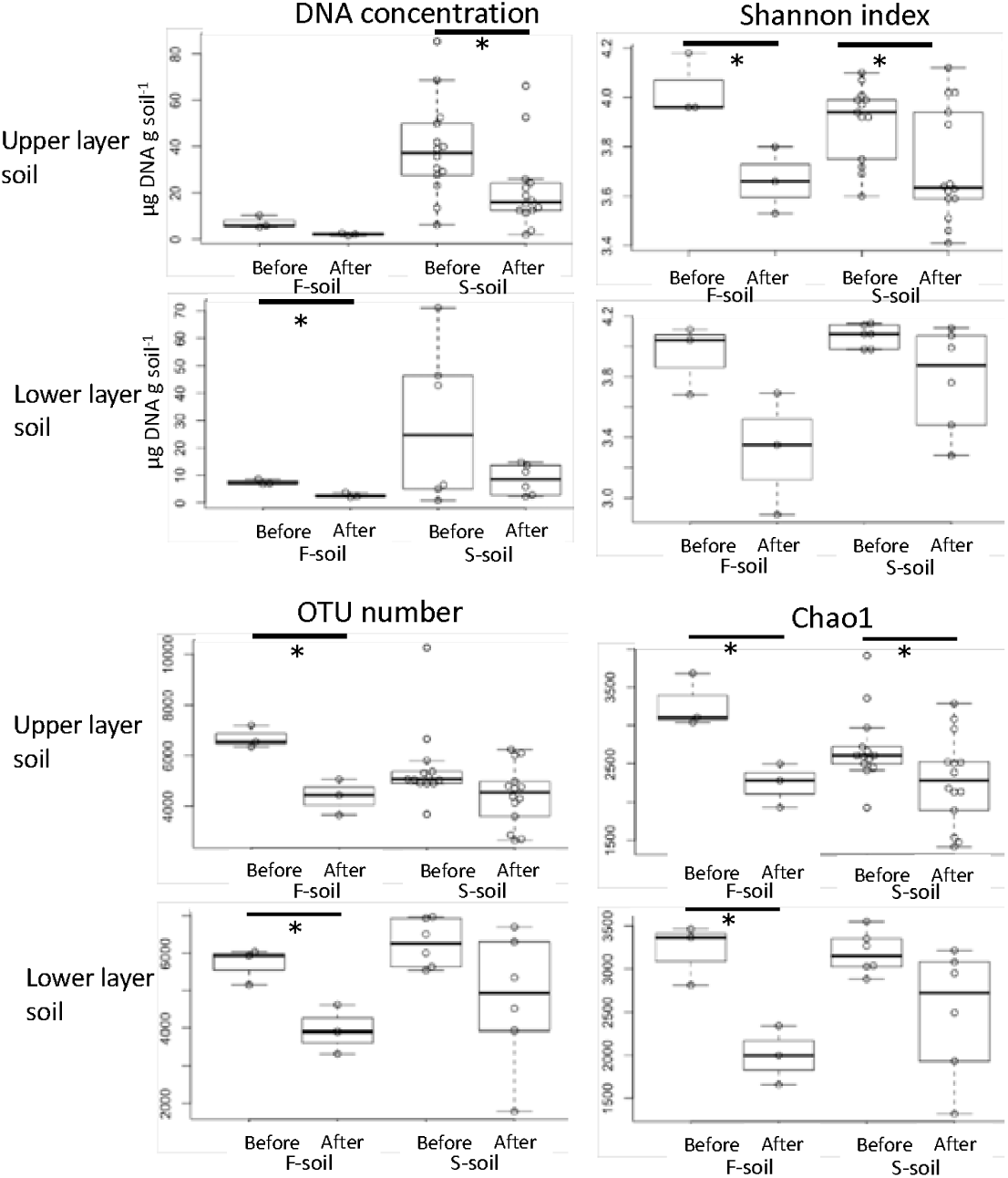
Comparison of DNA concentrations and prokaryotic diversity and richness between the F- and S-soils. Asterisks represent a significant difference in the mean values before and after ASD treatment (Tukey’s test, p < 0.05). F-soil: not well-disinfested soil, S-soil: well-disinfested soil

The Shannon index and Chao1 were significantly decreased in the S- and F-soils after ASD treatment of the upper layer. OTU numbers before and after soil sampling were significantly decreased in the F-soils regardless of the soil layer.

PCoAs based on weighted UniFrac analysis showed that the prokaryotic communities were roughly classified by sampling fields before and after ASD treatment (Figure 2).

**Figure 2.**
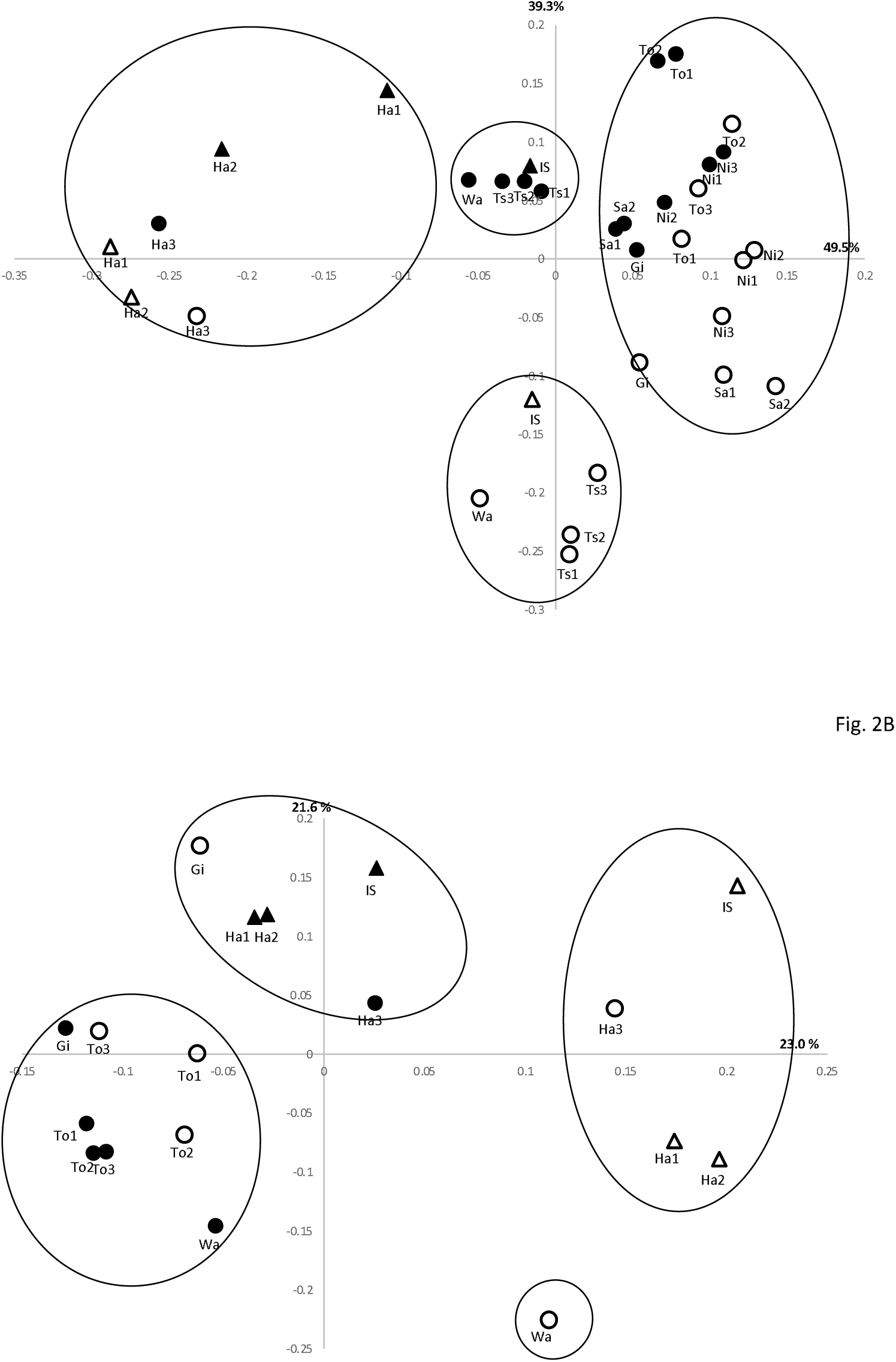
A UniFrac-weighted principal component analysis of the soil prokaryotic communities in the upper layer (A) and lower layer (B) of the field before and after ASD treatment. Closed circle: S-soil before ASD treatment, Open circle: S-soil after ASD treatment, Closed triangle: F-soil before ASD treatment, Open triangle: F-soil after ASD treatment. The clustering was conducted using K-means analysis.

The microbes collected from the same field comprised the same cluster before and after ASD treatment, despite the difference in ASD efficiency.

In the upper layer, field Ha contained one cluster regardless of ASD treatment and disinfestation efficiency (Figure 2A). Despite separation on the basis of before and after ASD treatment in fields Is, Wa, and Ts, the disinfestation efficiency had no effect. There were differences in the microbial communities of the lower layer soils of fields Is, Ha, and Gi after ASD treatment; however, no differences were detected between the S- and F-soils (Figure 2B). After ASD treatment, a more than two-fold increase in the relative abundance of Firmicutes was observed in eight greenhouses (Supplemental Table 3). In the upper layer soil, the relative abundance of Firmicutes was significantly increased after ASD treatment, whereas that of Crenarchaeota was decreased (Figure 3).

**Figure 3.**
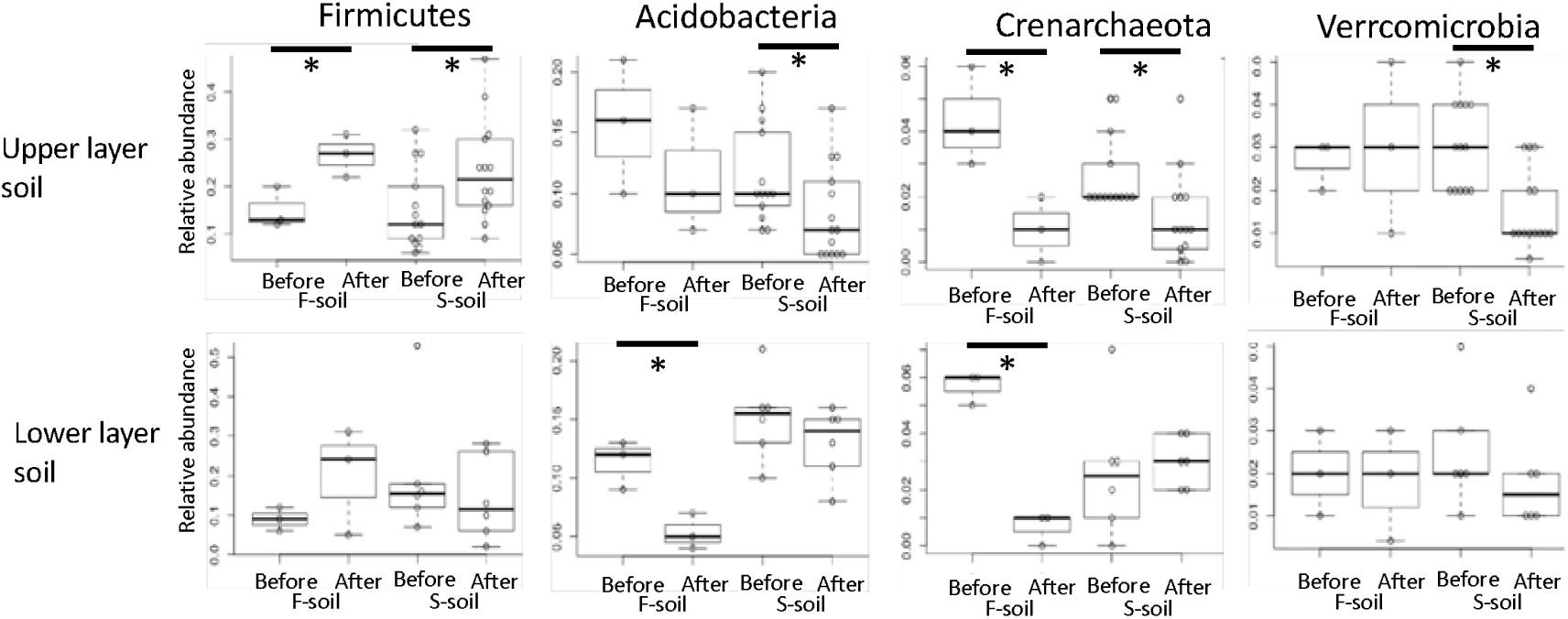
Comparison of the prokaryotic phyla with significant differences in abundance between the S- and F-soils. Asterisks represent a significant difference in the mean value of before and after ASD treatment (Tukey’s test, p < 0.05). F-soil: not well-disinfested soil, S-soil: well-disinfested soil

The relative abundance of *Acidobacteria* and *Verrucomicrobia* in the upper layer of the S-soil remained unchanged, whereas that of *Acidobacteria* and *Crenarchaeota* in the lower layer of the F-soil was significantly decreased. At the class level, a more than two-fold increase in the relative abundance of *Clostridia* of the phylum Firmicutes was observed after ASD treatment in the upper and lower layers of 15 (93.8%) of 16 and 8 (88.9%) of nine greenhouses, respectively (Supplemental Table 4). Moreover, the relative abundances of *Betaproteobacteria* and *Clostridia* were significantly increased after ASD treatment in the upper layer of the S-soil (Figure 4). In the S-soil, *Sphingobacteria* and *Cytophagia* were significantly decreased in the upper layer, and *Thermoleophilia* and *Actinobacteria* were significantly decreased in the lower layer.

**Figure 4.**
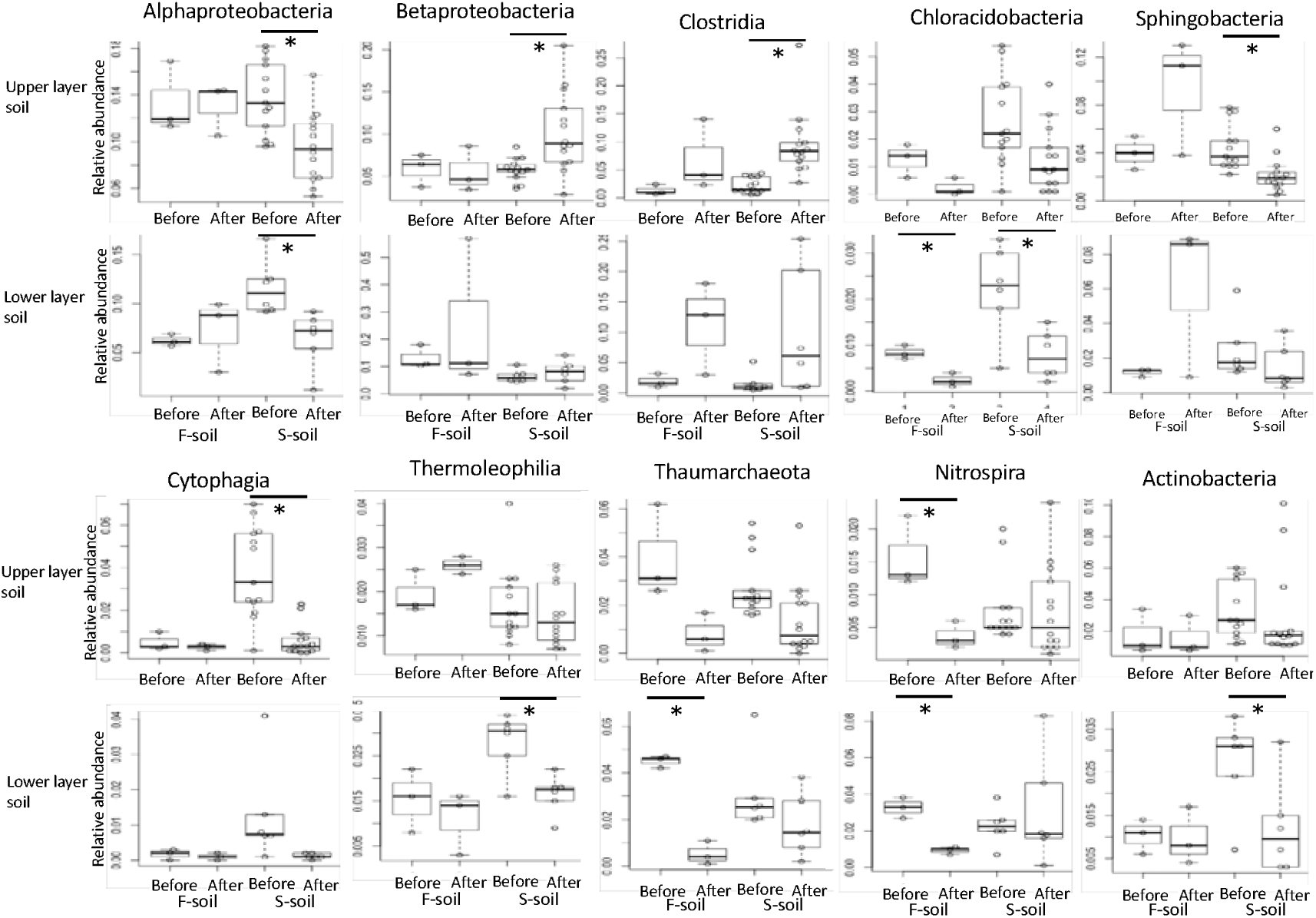
Comparison of the prokaryotic class with significant differences in abundance between the S- and F-soils. Asterisks represent a significant difference in the mean values before and after ASD treatment (Tukey’s test, p < 0.05). F-soil: not well-disinfested soil, S-soil: well-disinfested soil

### 3.4. Specific increased OTUs in various S-soils

Indicspecies analysis detected 25 OTUs from the upper and lower layers that were specifically increased in the S-soil after ASD treatment in >8 (50%) of 16 sites and 4 (66.7%) of 6 sites, respectively (Table 3).

**Table 3.**
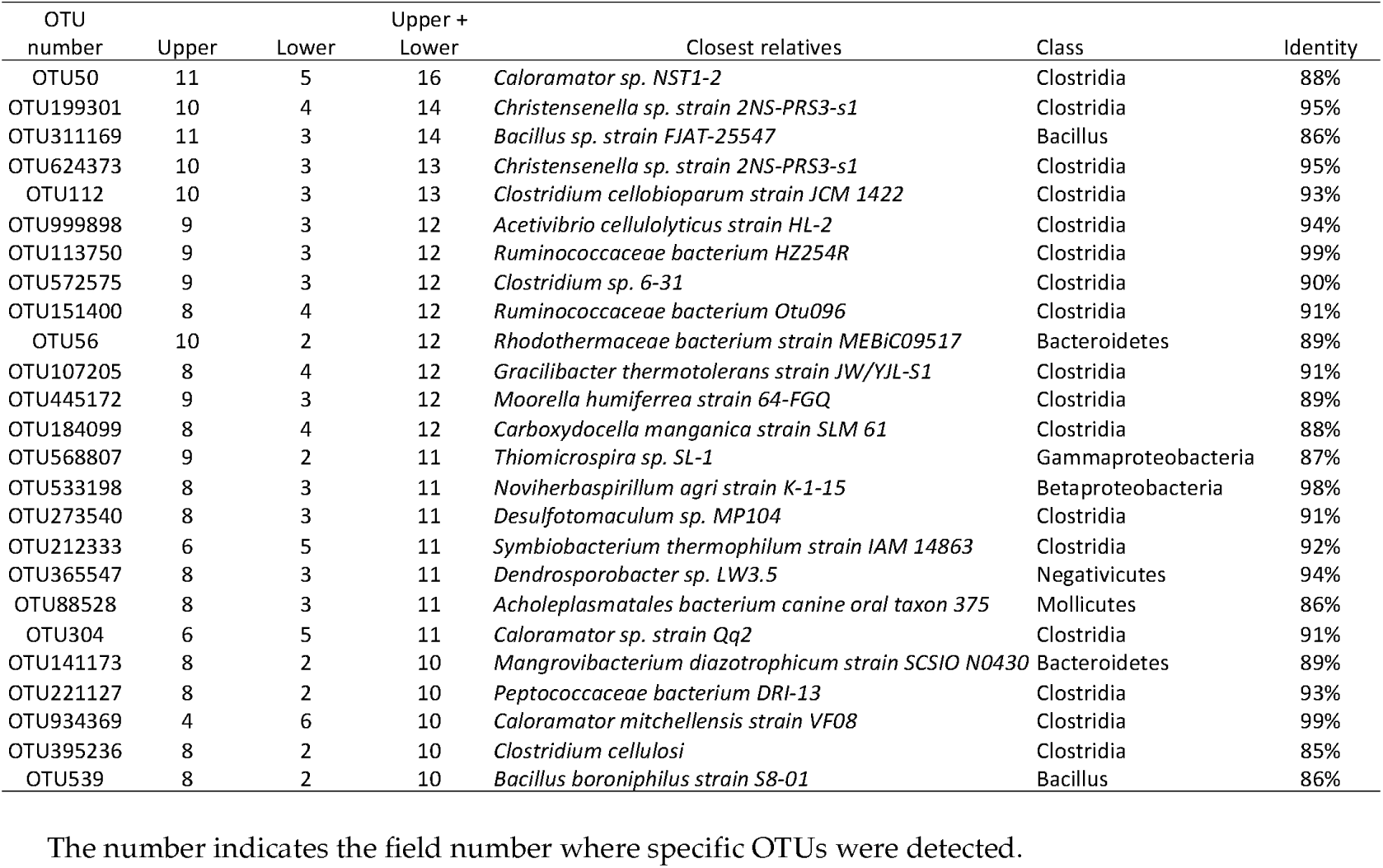
Shared prokaryotic OTUs with significantly increased abundance in the S-soils of different fields

The identified microbes belonged to several classes, including Clostridia, Bacillus, Betaproteobacteria, Gammaproteobacteria, and Mollicutes. Clostridia accounted for 18 (72%) of the 25 OTUs. In a phylogenetic tree of OTUs related to clostridial members, various Clostridia species were found to be specifically increased after ASD treatment in the S-soil (Figure 5).

**Figure 5.**
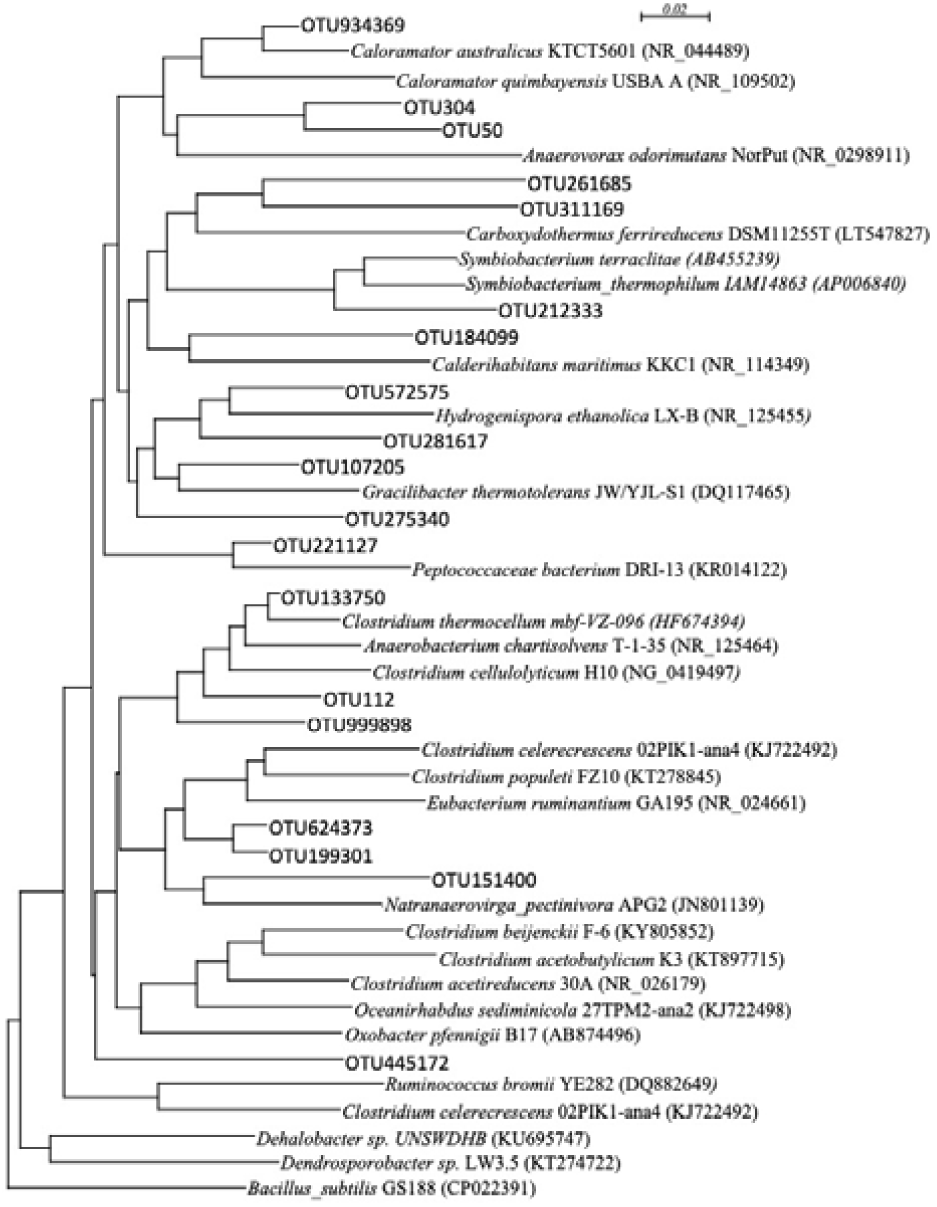
Neighbor-joining tree showing the phylogenetic relationships of the clostridial OTUs based on 16S rRNA gene sequencing (According to the clostridial cluster analysis [25]). OTUs obtained from this study are shown in gothic letters. Bacillus subtilis was used as the outgroup. Gothic letters indicate OTUs that specifically appeared in the S-soil.

Three OTUs belonged to the *Caloramator* group, *Clostridium* cluster III, whereas two OTUs belonged to the *Christensenella* group. OTU 50 (*Caloramator*) was commonly detected in 11 upper layer and 5 lower layer soil samples. Furthermore, OTU311169 (*Bacillus*), OTU199301 and OTU624373 (*Christensenella*), OTU 112 (*Clostridium*), and OTU56 (*Rhodothermaceae*) were increased in 10 (71.4%) of the 14 S-soils, whereas OTU934369 and OTU304 (*Caloramator*) and OTU212333 (*Symbiobacterium*) were commonly detected in >5 of the 6 fields of S-soils. On the other hand, no prokaryotes reduced by ASD were found to be common in more than half of the fields (data not shown).

## 4. Discussion

The results of the present study showed no significant differences in the abundances of microbial communities among fields with different disinfestation effects. In our previous study, prokaryotic communities were more affected by indigenous communities than the disease suppression effect [19]. The soil from a specific site typically contains microbial communities of greater similarity irrespective of soil treatment [26]. In the Ha field, three contiguous greenhouses received the same field management regimen for >10 years. Although there was a difference in the disinfestation effects, these samples were classified into the same cluster. These results suggest that the differences among indigenous soil communities are unaffected by controlling soil-borne pathogens.

Several OTUs that mainly belonged to *Clostridia* were specifically increased in several S-soils despite differences in indigenous prokaryotic communities. *Clostridia* are gram-positive anaerobic bacteria. Several studies have reported that the abundance of Clostridia species increases and it becomes the dominant bacteria after ASD treatment using plant material or low-concentration ethanol as a substrate [13,14,27]. During ASD, decreased oxygen utilization promotes the increased prevalence of anaerobic microbes [6,12,27]. Effective ASD requires sufficient soil reduction [28]. Therefore, well-disinfested soil is a good reduction condition, which may promote the growth of several anaerobic bacteria, such as *Clostridia*. Moreover, some *Clostridia* can produce spores that are resistant to high temperatures during the disinfestation period. Therefore, environmental changes might have promoted the increased abundance of Clostridia, some of which play important roles in the suppression of pathogen growth and proliferation. *Caloramator* are rod-shaped, obligate anaerobic, thermophilic endospore-producers that ferment several sugars and produce acetate, isobutyrate, lactate, and other volatile fatty acids (VFAs) but do not degrade cellulose [25,29]. More than half of clostridial clones that were increased after ASD treatment belonged to Clostridium cluster I, which included the *Oxobacter* and *Caloramator* groups [30]. *Symbiobacterium* spp. is a rod-shaped thermophiles that are syntrophic with *Bacillus* spp.; which possess a glucose degradation pathway [31]. *Christensenella* is a non-spore forming, short, straight rod with tapered ends and can use various sugars and produce VFAs during fermentation [32]. These results suggest that phylogenetically diverse Clostridia members might be responsible for the suppression of pathogens via the production of VFAs during the anaerobic decomposition of sugar-containing diatoms.

Moreover, other bacteria were increased after ASD in several S-soils. For example, *Noviherbaspirillum*, belonging to the class Betaproteobacteria, were specifically detected in the S-soils. *Noviherbaspirillum* are gram-negative, aerobic, non-spore forming rods capable of using several carbohydrates as carbon sources [33]. Betaproteobacteria increased after ASD treatment with wheat bran application in soil depths of 15.2 and 45.7 cm [34]. Moreover, several *Bacillus* species have been described as biological control agents against bacterial wilt and have been detected in soil after ASD treatment [30,35,36]. Cytophagaceae bacteria were significantly increased after ASD treatment with rice bran [16]. Although not previously detected in ASD-treated soil, *Symbiobacterium, Christensenella, Noviherbaspirillum*, and *Rhodothermaceae* might play important roles in the disinfestation of *R. solanacearum. Noviherbaspirillum, Bacillus*, and *Rhodothermaceae* are aerobic bacteria, but were increased by ASD, just as the anaerobic bacteria. In the soil environment, some *Bacillus* might contribute to the rapid decrease of the soil redox potential at the initial stage of ASD treatment via the consumption of oxygen [30]. Therefore, they might have important roles in maintaining anoxic soil conditions during ASD treatment. The abundances of Bacteroidetes, Acidobacteria, Planctomycetes, and Gammaproteobacteria increased after ASD treatment and some have been associated with the improvement of crop yields and suppression of plant diseases [26,37–39]. The fungi of the genus *Zopfiella*, which have been found to cause disease after ASD treatment, can prevent damping-off disease in cucumber [40]. Therefore, the aerobes detected in this study have the potential to directly suppress the growth and proliferation of plant pathogens.

The results of this study showed that *Clostridia* members were increased after effective ASD treatment in several fields. Moreover, some may be involved in the suppression of pathogen proliferation during ASD treatment. However, the killing mechanism of pathogens by ASD is not well -known. Some studies have shown that VFAs or ferrous iron can directly kill pathogens [15,41]. Future isolation studies are required to clarify the roles of the microbes detected in this study. These microbes may be involved in the suppression of pathogen proliferation; therefore, they could be good indicators of successful disinfestation by ASD. By revealing the ecology, there may be several strategies to increase the abundances of beneficial microbes.

In conclusion, prokaryotic communities were not strongly affected by the disinfestation efficiency among the sampling fields. However, the relative abundance of *Clostridia* and *Betaproteobacteria* were significantly increased in well-disinfested soil. Further, 25 OTUs were specifically detected in S-soils and most were affiliated to *Clostridia*, which is well-known to promote the effects of ASD. However, some microbes were not previously detected in ASD-treated fields. These microbes present good indicators of well-disinfested soil. Nonetheless, future studies are required to further elucidate the role of these microbes in disinfestation.

## Funding Information

This work was supported by the Cabinet Office, Government of Japan, Cross-ministerial Strategic Innovation Promotion Program (SIP), and the “Technologies for creating next-generation agriculture, forestry and fisheries” (funding agency: Bio-oriented Technology Research Advancement Institution, NARO). This work was also partially supported by the RIKEN Competitive Program for Creative Science and Technology, a Grant-in-Aid for Scientific Research from the Japan Society for Promotion of Science, No. 17K01447 (to M.O.), and by the Sasakawa Scientific Research Grant from The Japan Science Society (to C.G.L.). We would like to thank Editage (www.editage.jp) for English language editing.

### Supplementary Materials

The following are available online at www.mdpi.com/link, Table S1: Read numbers of the 16S rRNA gene in each soil sample, Table S2: The ratio of the DNA concentrations, prokaryotic diversity, and richness before to those after ASD treatment, Table S3: The ratio of the prokaryotic phylum before to that after ASD treatment, Table S4: The ratio of the prokaryotic class before to that after ASD treatment.

## Acknowledgments

We would like to thank Enago (www.enago.jp) for the English language editing.

## Author Contributions

All authors contributed to the intellectual input and provided assistance to this study. Each author contributed to conducting the ASD in each prefecture (E. M. and K. Y. in Ishikawa, M. K. in Toyama, M. M. in Niigata, Y. M. and H. W. in Gifu, and Y. O. in Wakayama prefecture). T. I., K. N., and M. O. have supervised the work and approved the manuscript for publication.

## Conflicts of Interest

The authors declare no conflicts of interest.

## Notes

#### Summary of Updates

The MS was conducted by English proofing.

