## Supplemental datasets for "Comparison of prokaryotic communities among fields exhibiting different disinfestation effects by anaerobic soil disinfestation"

Supplemental Table 1. Read numbers of the 16S rRNA genes in each sample

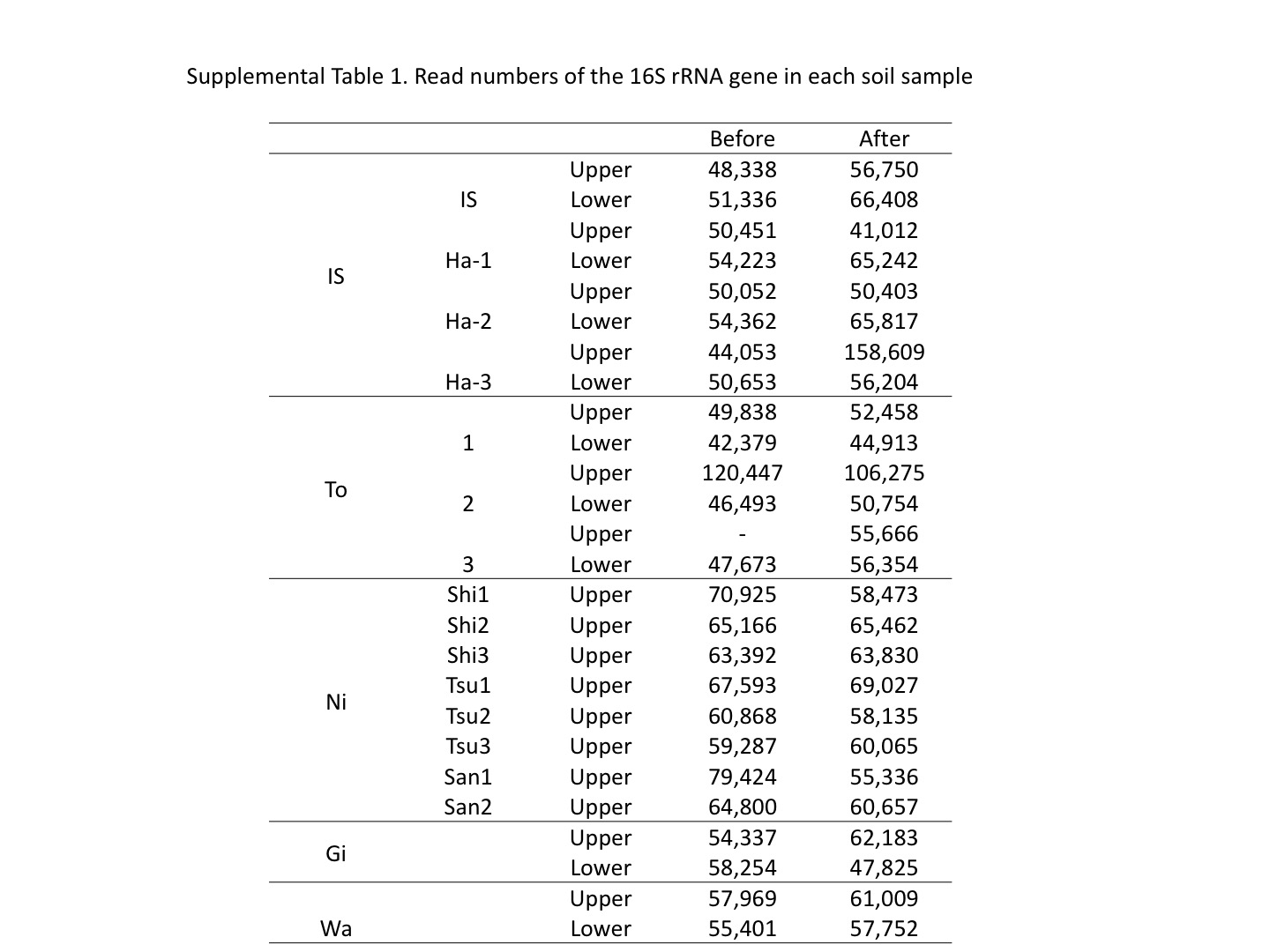

IS: Ishikawa field, Ha; Hakusan field, To; Toyama field, Ni; Niigata field, Tsu; Tsubame field, Sa; Sanjo field; Gi; Gifu field, Wa; Wakayama field. The numbers after each field indicates the site number.

Supplemental table 2. The ratio of the DNA concentration, prokaryotic diversity and richness before to those after ASD treatment

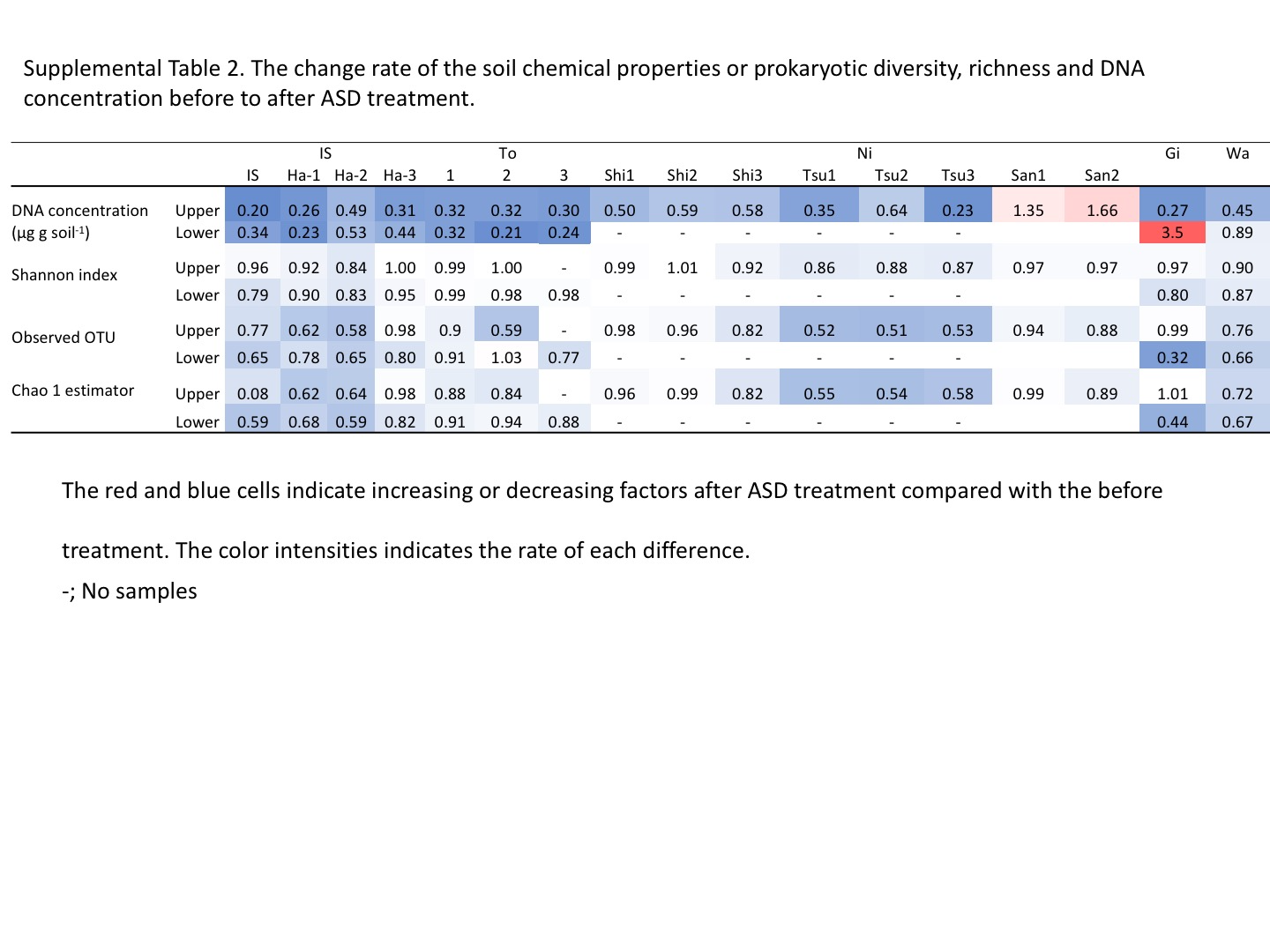
The red cells indicate increasing factors after ASD treatment when compared with the before treatment. The blue cells indicate decreasing factors after ASD treatment when compared with the before treatment. The color intensities indicate the ratio of each difference.

-; No samples

Supplemental table 3. The ratio of the prokaryotic phylum before to that after ASD treatment

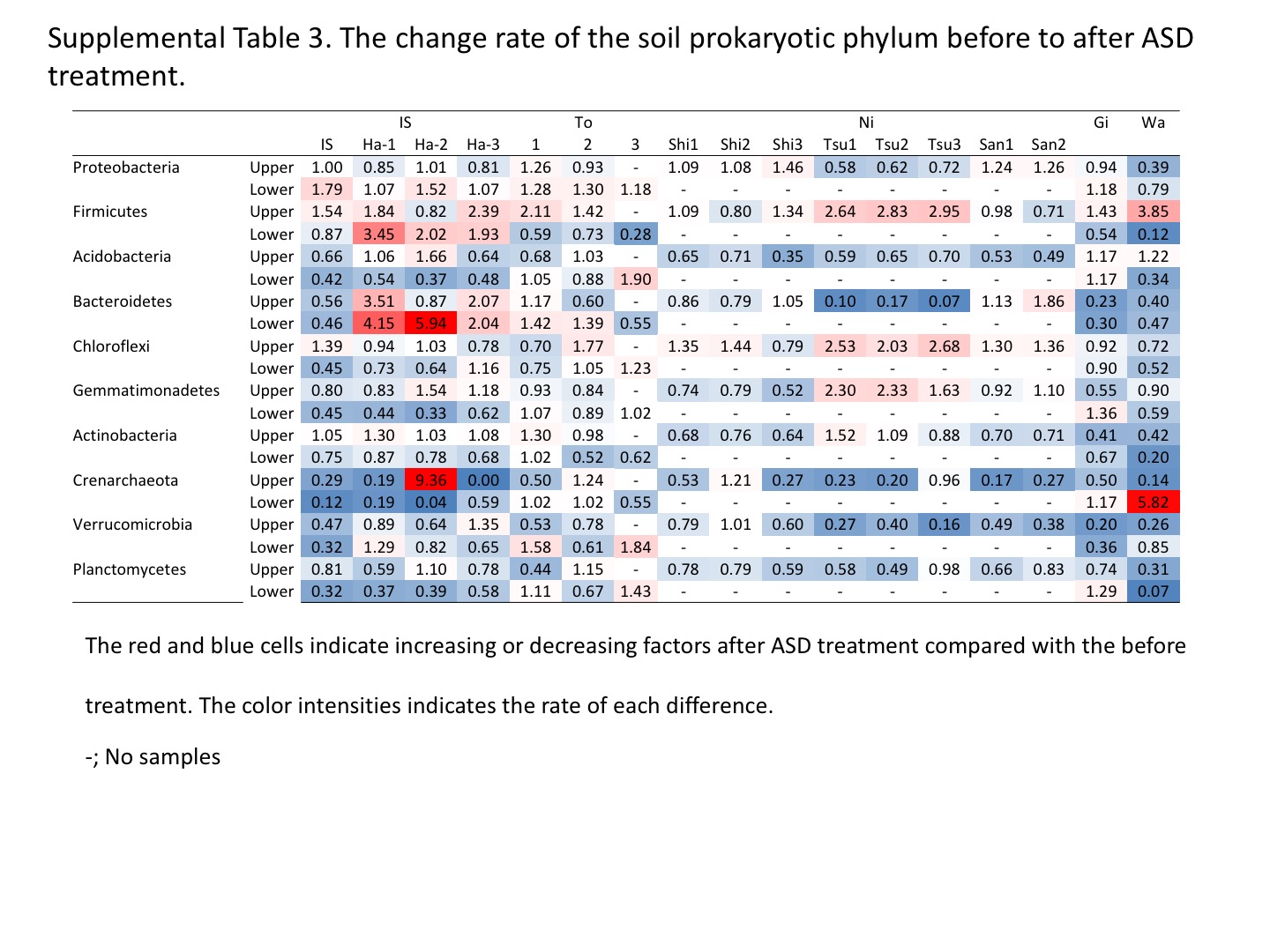

The red cells indicate increasing factors after ASD treatment when compared with the before treatment. The blue cells indicate decreasing factors after ASD treatment when compared with the before treatment. The color intensities indicate the ratio of each difference.

-; No samples
Supplemental table 4. The ratio of the prokaryotic class before to that after ASD treatment

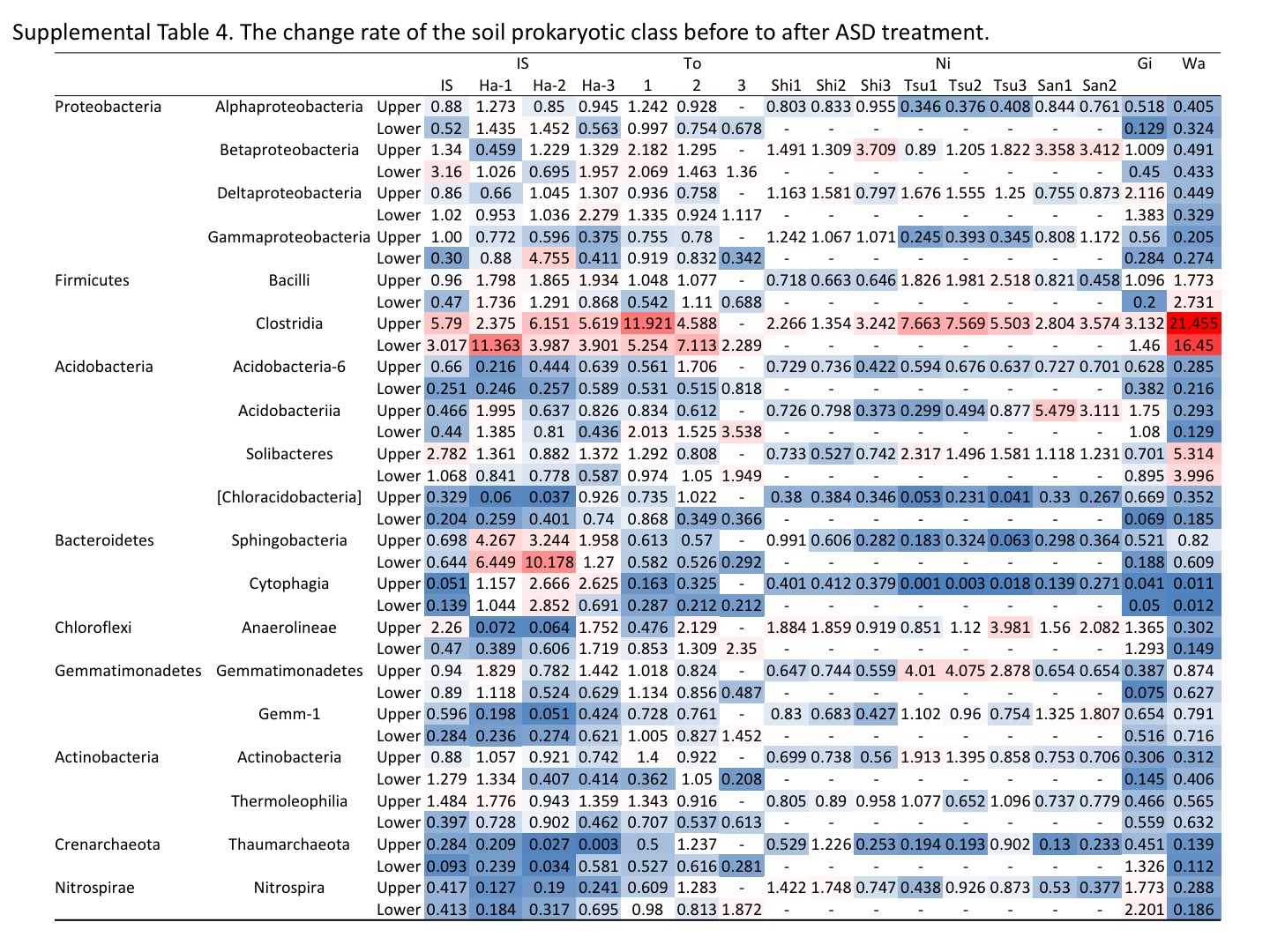

The red cells indicate increasing factors after ASD treatment when compared with the before treatment. The blue cells indicate decreasing factors after ASD treatment when compared with the before treatment. The color intensities indicate the ratio of each difference.

-; No samples
